# InDel assembly: A novel framework for engineering protein loops through length and compositional variation

**DOI:** 10.1101/127829

**Authors:** Pedro A. G. Tizei, Emma Harris, Marleen Renders, Vitor B. Pinheiro

## Abstract

Insertions and deletions (indels) are known to affect function, biophysical properties and substrate specificity of enzymes, and they play a central role in evolution. Despite such clear significance, this class of mutation remains an underexploited tool in protein engineering with no available platforms capable of systematically generating or analysing libraries of varying sequence composition and length. We present a novel DNA assembly platform (InDel assembly), based on cycles of endonuclease restriction and ligation of standardised dsDNA building blocks, that can generate libraries exploring both composition and sequence length variation. In addition, we developed a framework to analyse the output of selection from InDel-generated libraries, combining next generation sequencing and alignment-free strategies for sequence analysis. We demonstrate the approach by engineering the well-characterized TEM-1 β-lactamase Ω-loop, involved in substrate specificity, identifying multiple novel extended spectrum β-lactamases with loops of modified length and composition areas of the sequence space not previously explored. Together, the InDel assembly and analysis platforms provide an efficient route to engineer protein loops or linkers where sequence length and composition are both essential functional parameters.

## Introduction

Directed evolution is a powerful tool for optimizing, altering or isolating novel function in proteins and nucleic acids^1,2^. Cycles of sequence diversification to generate libraries followed by partitioning of those populations through selection, enable a desired function to be isolated and systematically optimised. Directed evolution is therefore a walk across sequence space with library quality and diversity as key factors in how efficiently that search can be carried out and on how much of the available sequence space can be explored.

Current library synthesis methods that exploit compositional variation focus on generating libraries of constant length, capable of efficiently sampling a given sequence landscape (of fixed-length) but unable to explore the entire available sequence space, i.e. landscapes of different lengths. They vary in cost, in how that diversity is distributed and in the level of customization (i.e. redundancies, biases and coverage) that can be implemented^3,4^. PCR based methods using degenerate primers provide a cost-effective route towards creating focused (i.e. that target a small number of clustered sites) high-quality libraries^5^ but cannot rival commercial high-throughput DNA synthesis platforms^6^, or DNA assembly methods that rely on the incorporation of individual triplets^7,8^, for customization.

Methods have been developed to exploit changes in length – be it through modifying oligo synthesis^9^, by using insertion and excision cycles of engineered transposons^10^ or by combining chemical and enzymatic approaches^11^ – can deliver high quality libraries (i.e. where most indels do not affect the reading frame) but need specialist equipment or are only able to generate a limited spectrum of indel mutations.

This is particularly relevant to the engineering of systems in which loops contribute significantly to protein function, such as the H3 loop in antibodies^12^ or the loops in TIM barrel enzymes^13^. In those circumstances, methods that target loop composition as well as length are essential for efficient functional optimization. Traditional methods can address the problem by brute force, sampling sequence space through the use of multiple libraries of varying sequence composition, each with a given length^14^. Nonetheless, the challenge for analysing the output of selection of such library remains unaddressed.

Selection can be carried out until population diversity is sufficiently low that analysis is redundant, or by analyzing single-length landscapes^15^. The first approach increases the possibility of failure (e.g. parasites in selection), and can lead to the isolation of suboptimal variants because of experimental biases and inadequate sampling in the early selection rounds. Deep sequencing of the libraries captures the complexity of the available functional space in early rounds but, by analysing single-length landscapes individually, some information is inevitably lost, potentially by masking motifs present in multiple lengths or by increasing the possibility of false positives in sparsely sampled landscapes.

Here, we present a combination of (i) a cost-effective DNA assembly of high-quality, highly customizable focused libraries capable of sampling both length and compositional variation, and (ii) a robust analytical framework that utilizes deep sequencing of pre- and post-selection libraries to identify enriched motifs across different length libraries. Together, they establish a powerful strategy to efficiently engineer loops and linkers from repertoires that vary in both length and composition.

## Results

### DNA assembly by cycles of restriction and ligation – the InDel assembly

InDel assembly relies on cycles of DNA restriction and ligation to progressively assemble a DNA library on paramagnetic beads, which serve as solid support and facilitate sample handling. The assembly starts with a biotinylated dsDNA template, encoding the starting point of the library and a recognition site for a type IIs endonuclease, bound to paramagnetic streptavidin-coated beads (Fig. 1a).

**Figure. 1.**
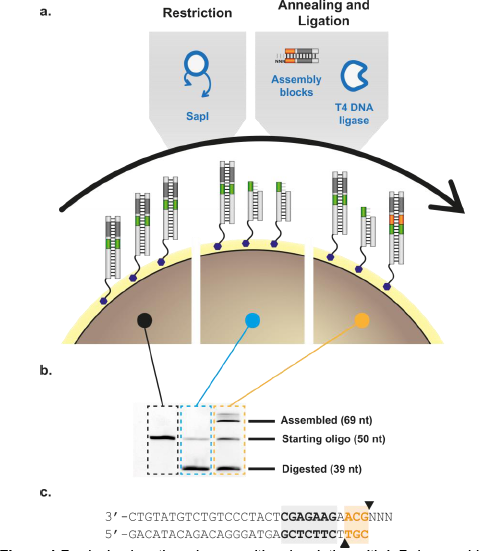
Producing length and compositional variation with InDel assembly. (a) At each assembly cycle, dsDNA templates bound to the paramagnetic beads are restricted with SapI (a type IIs endonuclease), building blocks annealed and ligated. After ligation, the cycle can be restarted. Compositional variation is achieved primarily by combining pools of different building blocks. (b) Denaturing gel electrophoresis of fluorescently labelled template across the different steps of the cycle show that restriction and ligation are not carried out to completion in any step, underpinning the length variation of the resulting libraries. (c) Sequence of a building block. A long double-stranded region is used to stabilize building block annealing and ensure efficient endonuclease activity. SapI recognition site is shown in grey and restriction sites as black triangles. The overhang depicted would add GCA, coding for alanine to the growing chain ‒ further information in Supplementary Table 2.

Type IIs restriction endonucleases bind non-palindromic recognition sequences and cleave dsDNA specifically, generating blunt or short single-stranded overhangs, which have been extensively exploited in molecular biology for ‘seamless’ cloning, as in Golden Gate assembly^16^. In InDel assembly, SapI (a type IIs endonuclease) is used to digest the template, generating a 3-base ssDNA overhang and removing its recognition site from the bead-bound template.

The overhang generated by that cleavage enables the ligation of a standardized dsDNA building block with compatible overhangs. Building blocks (Fig. 1c) have been designed to have a degenerate overhang (minimising template sequence constraints) and a SapI site, which enables the assembly cycle to be restarted.

Crucially, a triplet is placed between overhang and SapI cleavage site, ensuring an increase in sequence length and maintaining the underlying reading frame. Variation of the triplet, which can be achieved by concomitantly adding two or more building blocks (in practice, any custom mixture thereof) to the ligation step, therefore leads to sequence variation in the resulting library. Like Sloning^8^ and ProxiMAX^7^, InDel is capable of delivering a highly flexible library since the building blocks can be mixed in any ratio and can incorporate any sequence and length of DNA – and hence could also be explored for protein fragment assembly.

Because no restriction or ligation reaction is carried out to completion in the system (Fig. 1a and Fig. 1b), the library accumulates not only compositional but also length variation with a fraction of available templates not extended in each assembly cycle. The resulting InDel assembled library therefore is more complex than what can be achieved by commercial platforms – it generates diversity comparable to COBARDE^9^ but requires no specialist equipment or reagents for library assembly.

Each reaction step in the InDel assembly was validated and optimised using fluorescently labelled templates, with restriction and ligation monitored by shifts in mobility of the fluorescent oligo in denaturing PAGE (Fig. 1b). We optimized ligation (Supplementary Fig. 1) and restriction conditions, explored building block topology (i.e. hairpins or dsDNA from annealed strands), explored creating library degeneracy through concomitant addition of multiple blocks, and other reaction parameters.

Optimised reactions suggested 50% assembly efficiency per cycle could be obtained, however later sequence analysis of synthesized libraries determined that the incorporation efficiency per cycle was lower ‒ probably the result of extended sample handling and the limited activity of SapI in extended reactions. Further optimization of assembly conditions, in the form of the method presented here, yielded assembly efficiencies close to 30% per cycle (Supplementary Fig. 2). Codon biases were observed but varied between assembled libraries, suggesting that it is not a limiting factor in assembly and, as with similar platforms, can be further optimised if needed^7^.

### TEM-1 Ω-loop functional sequence space includes loops of different length as well as different composition

Having established the assembly platform, we chose the β-lactamase TEM-1 to demonstrate its potential. TEM-1 is a well-characterized enzyme^17,18^ that due to its ease of selection, wide range of available substrate analogues and its clinical relevance, has long been used as a model in directed evolution^19,20^. In particular, TEM-1 contains a short flexible loop which is part of its active site (the Ω-loop, _164_RWEPE_168_ ‒ Fig. 2a) and has been implicated in substrate specificity. To date, engineering of the Ω-loop has focused exclusively on exploring variation in composition of the loop, culminating on the successful isolation of _164_RYYGE_168_, a variant resistant to the cephem ceftazidime^21,22^.

**Figure 2.**
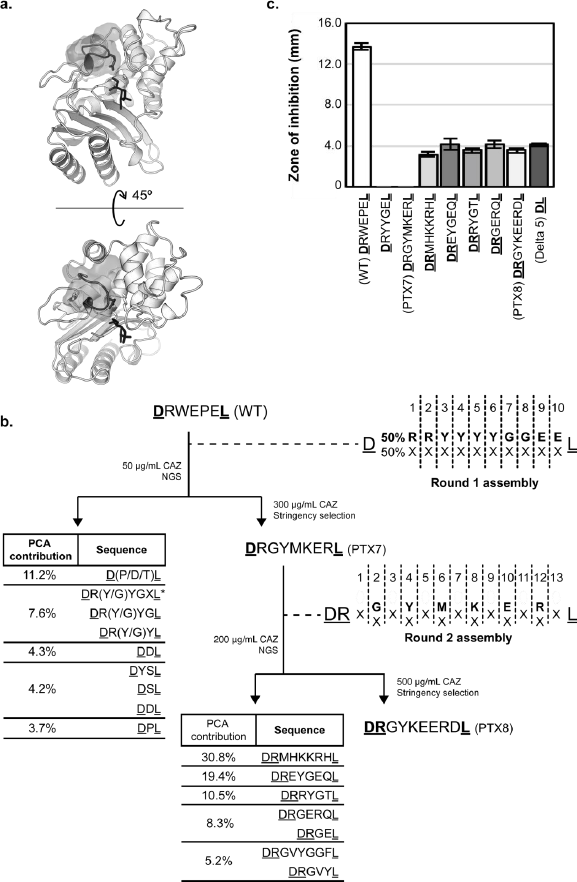
Directed evolution of TEM-1 Ω-loop variants with altered substrate specificity. (a) TEM-1 beta-lactamase in complex to substrate analogue inhibitor (PDB: 1TEM). Inhibitor is shown in the TEM-1 active site in black, and side chains of key catalytic residues (S70, K73 and E166) are shown. The Ω-loop is shown in grey and superimposed to its translucent space-filling representation. (b) Summary of the directed evolution of TEM-1. A 1st round library, biased towards the generation of RYYGE variant was assembled and selected. Next generation sequencing (NGS) confirms RYYGE was significantly enriched (DRYYGEL was the 16th most frequent sequence post-selection and picked up in the 2nd PCA dimension ‒ see Supplementary Table 1). PTX7, isolated from a high-stringency selection, was used as seed for the 2nd round library. PTX8 and sequences from the top four PCA dimensions were further characterized. (c) Ceftazidime resistance of selected variants, measured by inhibition of growth around antibiotic-soaked paper discs ‒ higher values indicate lower resistance (n = 3, error bars represent s.e.m). Underlined residues represent the invariant edges of the assembly, constant in each library and required for library amplification.

Our initial goal was to explore the sequence neighbourhood of the previously reported _164_RYYGE_168_ variant with a view towards demonstrating assembly and selection. Based on our early estimates of 50% assembly efficiency per cycle (Fig. 1b), we assembled a 10-cycle InDel library using biased mixes of building blocks ‒ 50% coding for the desired target residue and the remaining 50% divided between the remaining 19 coding triplets (Fig. 2b). Based on a simple binomial model, the library was expected to fully sample all variants of up to four inserted codons and sample longer landscapes increasingly more sparsely ‒ but always biased towards sequences related to _164_RYYGE_168_ (Supplementary Fig. 3).

Assembled libraries were cloned into a TEM-1 backbone harbouring the M182T stabilizing mutation^20^. Selection was carried out by plating cells transformed with the TEM-1 library directly on media supplemented with 50 μg/mL ceftazidime ‒ below the minimal inhibitory concentration for the _164_RYYGE_168_ variant harbouring the stabilizing mutation (Supplementary Fig. 4).

Sequencing of pre- and post-selection library confirmed that the _164_RYYGE_168_ variant was present in the starting library, albeit at a frequency lower than expected (5 reads in 2.3 million ‒ 0.0002% of the population), and was significantly enriched on selection (314 reads in 230,000 ‒ 0.14%) ‒ an enrichment score of 8460 (in the 99^th^ percentile of a distribution of enrichment Z-scores based on the comparison of two Poisson distributions). Enrichment of the TEM-1 _164_RYYGE_168_ variant in selection clearly demonstrates that InDel assembly and selection can recapitulate previous engineering efforts at altering the substrate specificity of TEM-1.

In parallel with deep sequencing of the libraries, we also increased the stringency of selection by plating the library at higher ceftazidime concentrations. At ceftazidime concentrations of 300 μg/mL, a single variant was isolated: _164_RGYMKER_168b_ (adopting antibody annotation to describe insertions^23^ ‒ see materials and methods for more details), differing in both composition and length from wild-type (_164_RWEPE_168_) and engineered TEM-1 (_164_RYYGE_168_) sequences. Undetected in the input library, _164_RGYMKER_168b_ represented approximately 0.004% of the selected library (9 reads in 230,000) and displayed a resistance profile comparable to that of the previously engineered _164_RYYGE_168_ (Fig. 2c). Isolation of _164_RGYMKER_168b_ TEM-1 variant confirms that high levels of ceftazidime resistance are not unique to _164_RYYGE_168_ and further validate that loop length is a crucial parameter in protein engineering.

### InDel-assembled libraries are high quality and efficiently sample sequence landscapes of different lengths

In addition to enabling us to look at the impact of selection, deep sequencing of pre- and post¬selection InDel-assembled libraries allowed the quality of the libraries to be assessed, including biases, coverage and assembly errors (e.g. frameshifts).

The pre-selection libraries had the expected biases introduced in assembly (Fig. 2b), with preferred codons being overrepresented in the library – e.g. R and Y in the round 1 library (Supplementary Fig. 2). Pre-selection sequence diversity was high, with only the first R incorporation showing significant conservation (Fig. 3c), and showed complete or heavily biased coverage towards the target _164_RYYGE_168_ motif (Fig. 3b). Post-selection, consensus motifs can be determined for each single-length sequence landscape but are strongest in the 5-mer landscape as RXYGX (matching to the previously described RYYGE motif), seen both as an increase in information content (Fig. 3d) and in the distribution of selected sequences across the landscape (Fig. 3e). This further confirms _164_RYYGE_168_ as a functional ‘peak’ in the 5-mer landscape.

**Figure 3.**
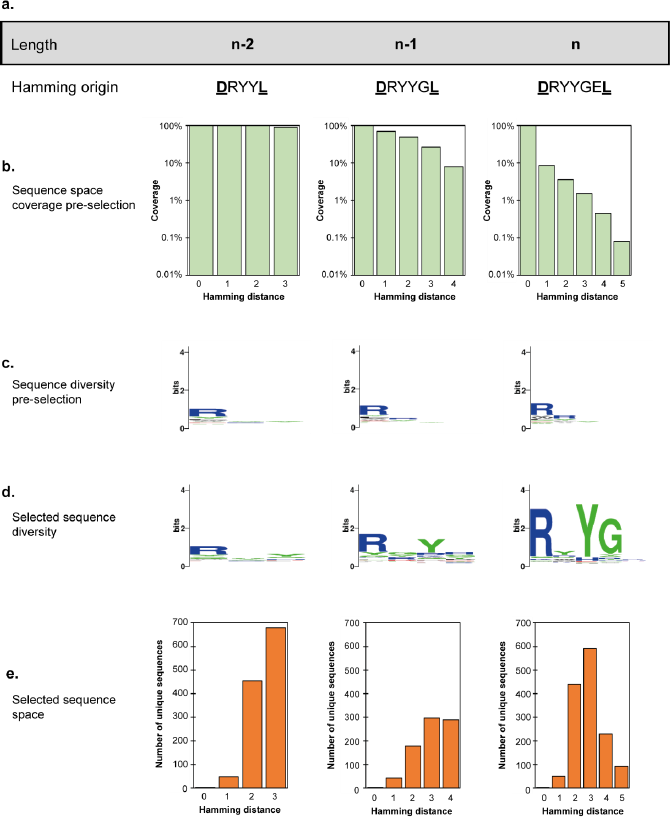
InDel assembly coverage of sequence space neighbouring RYYGE and impact of selection. (a) The available sequence space is split into fixed-length landscapes and each analysed separately using the most frequent variant of the desired length as the origin for Hamming distances. (b) The biased synthesis used in the InDel assembly of this first library ensured the sequence neighbourhood of the target RYYGE sequence was efficiently explored with lower Hamming distances more effectively sampled. (c) The library remains diverse with only minimal bias for arginine incorporation in the first position, as predicted. The height of each residue in the logo is a measure of their frequency at that position. (d) Selection clearly enriches for an RXY motif particularly in *n-1* and *n* landscapes. (e) Hamming distances to other unique sequences obtained in each landscape after selection, highlighting the presence of a local peak around RYYGE.

Analysis of enrichment also suggests that the functional space of the TEM-1 Ω-loop is densely populated, with multiple functional motifs present in different loop lengths – and not necessarily related to the wild-type _164_RWEPE_168_ or engineered _164_RYYGE_168_ motifs (Fig. 2c, Fig. 3 and Supplementary Fig. 5). This is further supported by our isolation of the _164_RGYMKER_168b_ variant, which differs from all previously reported variants in both composition and length, highlighting the power of InDel to navigate the sequence space in which length is an additional design parameter.

### Alignment-free sequence analysis improves identification of enriched motifs

While the use of deep sequencing to map functional landscapes^24^ and to accelerate directed evolution^25^ is well-established, current methods do not perform well for short libraries that vary in both length and composition^26^. Stratifying the library into fixed-length repertoires for analysis^15^ or using indels to contribute to a mismatch score (i.e. Hamming distance) have been applied to the analysis of length and compositional variation. However, length variation in directed evolution is generally discarded^27^ because of difficulties in positioning gaps in the resulting alignments^26^.

We therefore set out to develop an alignment-free sequence analysis method based on subdivision of sequence strings into short reading windows (*k*-mers) to extract information from sequencing data spanning multiple fixed-length sequence landscapes. *K*-mer based methods are integral components of large sequence comparison methods^28^ as well as next generation sequence assembly^29^, allowing comparison of sequences of different lengths as well as reconstruction of sequence motifs ‒ the two steps required to identify functional variants from the available InDel assembly libraries.

We opted for using masked 3-mers (i.e. X_1_X_2_X_3_ being considered as X_1_X_2__ and X_1__X_3_)^30^ to analyze sequences, reducing computational burden (the possible 8000 3-mers are reduced to 800 possible masked 3-mers) without significant loss of information relevant for motif reconstruction. Based on analysis of ‘toy’ data sets (not shown), we chose to explicitly represent residues flanking the synthesized libraries (as Z_1_ and Z_2_ ‒ see Fig. 4a) in analysis, adding a further 82 possible *k*-mers but significantly improving the robustness of downstream motif reassembly.

**Figure 4.**
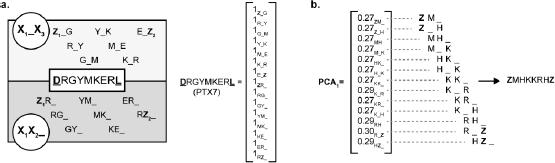
*K*-mer sequence decomposition and reconstruction. (a) Sequences can be decomposed into all possible masked 3-mers (i.e. X_**1**_X_**2**_X_**3**_ separated into X_**1**_X_**2**__ and X_**1**__X_**3**_) as shown for PTX7. Each masked 3-mer is counted generating an 882-dimension vector (non-zero elements shown). Vectors are normalized (multiplied by *V*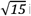 in the case of PTX7) and multiplied by their enrichment score. Principal component analysis (PCA) identifies enriched *k*-mers, which allow a sequence to be reconstructed. An arbitrary cut-off (0.1) can be used to minimize noise and facilitate assembly. (b) The sequence reconstruction of the first PCA component calculated from the second library selection.

Pre- and post-selection libraries were combined and each individual sequence, described by its masked 3-mer count, was treated as a column vector of 882 dimensions – the number of masked 3-mers used to describe sequence and library boundaries (Fig. 4a). Vectors were normalized and scaled by their Z-score as a measure of enrichment and as a proxy for function. Together, the entire output of selection is mapped onto a complex 882-dimensional space.

Principal component analysis (PCA) enabled us to deconvolute this complex space to identify, which combinations of the 882 dimensions (i.e. masked 3-mers) contribute the most to the distribution of the sequences within that space – in practice, allowing functional motifs to be reconstructed along individual PCA dimensions.

Highly enriched sequences contribute significantly to library variation and are identified in the first PCA dimensions (which account for the biggest variation in the data). Crucially, functional sequences that are related (i.e. share masked 3-mers) but not necessarily of the same length, cluster in this 882-dimensional space and are more readily picked up by analysis ‑ Table 1 and Supplementary Table 1.

Therefore, our approach identifies not only functional peaks that are restricted to a single length but also peaks that span different lengths, while considering contribution of submotifs (in longer loops) and degeneracies (i.e. non-conserved positions in a motif) that may also be contributing to selection. Motif reconstruction can be automated, with resulting sequences being tested for function or used as starting points for new libraries.

**Table 1.** Comparison of PCA-derived enriched sequences and NGS read frequency for the 2nd round library. Motifs were reconstructed for the first 10 PCA dimensions and used to search the NGS results for the ranking of the highest enriched sequences (highest Z-scores). Because the second round post-selection library was significantly smaller and heavily biased, correlation between PCA and NGS frequency is very good. Some sequence variation and small motifs can still be seen.

| PCA dimension | PCA-derived motif | Most frequent match | Frequency Ranking |
| --- | --- | --- | --- |
| 1 | <u>ZMHKKRHZ</u> | <u>ZMHKKRHZ</u> | 1 |
| 2 | <u>ZEYGEQZ</u> | <u>ZEYGEQZ</u> | 2 |
| 3 | <u>ZRYGTZ</u> | <u>ZRYGTZ</u> | 3 |
| 4 | <u>ZGERQZ</u><br><u>ZGEZ</u> | <u>ZGERQZ</u> | 4 |
| 5 | <u>ZGVYGGFZ</u> | <u>ZGVYGGFZ</u> | 7 |
|  | <u>ZGVYZ</u> | <u>ZGVYZ</u> | 8 |
| 6 | <u>ZAKERHZ</u> | <u>ZAKERHZ</u> | 5 |
|  | GVY | <u>ZGVYZ</u> | 8 |
|  | <u>ZE(V/K/R)XZ</u> | <u>ZEKERHZ</u> | 108 |
| 7 | <u>ZAKEXH</u> | <u>ZAKERHZ</u> | 5 |
|  | <u>Z(A/G)YVZ</u> | <u>ZGYVZ</u> | 6 |
| 8 | <u>ZEEVHZ</u> | <u>ZEEVHZ</u> | 9 |
| 9 | <u>Z(A/W)(E/Y)EHRZ</u> | <u>ZWEGRQZ</u> | 10 |
|  | <u>ZWEGR(Q)Z</u> | <u>ZAYEHRZ</u> | 12 |
|  | VEGRQ |  |  |
| 10 | <u>VXZ</u> |  |  |
|  | <u>Z(A/G)Y(Y/E)HRZ</u> | <u>ZGVYZ</u> | 8 |
|  | <u>ZGY(Y)Z</u> | <u>ZAYEHRZ</u> | 12 |
|  | <u>ZGAYEHRZ</u> |  |  |

### The sequence neighbourhood of TEM-1 Ω-loop is densely populated with functional variants

Sequence analysis of the first round of selection identified a wide range of sequence motifs that were enriched in selection (Supplementary Table 1), suggesting that the sequence neighbourhood of the Ω-loop was more densely populated than previously expected. The short motifs identified however could be a reflection of the lower assembly efficiency of the first round (biasing the Ω-loop library towards short motifs) as well as the low stringency of selection used (enabling even moderately active variants to be selected).

We therefore decided to pursue a second round of selection to investigate the sequence space in the neighbourhood of _164_RGYMKER_168b,_ including deletions, substitutions and insertions. Exploring that sequence space could easily be achieved with InDel by assembling a library alternating between fully degenerate (i.e. equal distribution of all available triplets) with biased (i.e. 50% of desired _164_RGYMKER_168b_ triplet and 50% of the remaining 19 triplets) cycles (Fig. 2b).

As before, selection was carried out by plating the assembled library onto solid media containing ceftazidime. Higher antibiotic concentrations were used (200 μg/mL) to increase selection stringency and 79 colonies were isolated. Pre- and post-selection libraries were sequenced and analysed with our *k*-mer approach (Table 1).

The top four candidates from the second-round library (**DR**MHKKRHL, **DR**EYGEQL, **DR**RYGTL, **DR**GERQL), harbouring five to eight amino acids in the diversified region of the Ω-loop, were further characterized (Fig. 2c and Supplementary Fig. 4). All four variants (as well as a shortened loop variant Δ5) are significantly more resistant to ceftazidime than the wild-type enzyme.

Characterised variants show little sequence similarity to wild-type or engineered variants (_164_RYYGE_168_ and _164_RGYMKER_168b_). This diversity confirms that the sequence neighbourhood of the Ω-loop is densely populated with functional variants in multiple landscapes. It also demonstrates the potential of the InDel framework to efficiently explore sequence space varying both length and composition.

## Discussion

Our results provide further evidence that loops are highly evolvable^31^ and also highlight how directed evolution of protein loops must take into account sequence spaces that straddle more than a single-length landscape. We show that the combination of InDel assembly and *k*-mer-based analysis provide a powerful framework for navigating sequence space that is not otherwise accessible. Effectively, InDel assembly, selection and *k*-mer analysis respectively provide ‘build’, ‘test’ and ‘learn’ steps of the Synthetic Biology cycle^32^ and could be automated to accelerate engineering of any protein function.

In addition, we present here an example of InDel assembly with triplets, which is ideal for generating amino-acid-steps in libraries of protein coding genes. The platform is compatible with building blocks of mixed length, enabling a vast host of combinatorial possibilities that could be applied to the directed evolution of nucleic acid aptamers, gene expression regulatory elements and fragment-based protein engineering.

## Acknowledgements

PAGT acknowledges CAPES foundation support (fellowship BEX 8985-13-8). VBP acknowledge support by the European Research Council [ERC-2013-StG project 336936 (HNAepisome)] and by the BBSRC (grant BB/K018132/1). The authors also thank Dr. Chris Cozens and Dr. Andrew Osbourne for critical reading of the manuscript.

## Author contributions

PAGT and VBP conceived the assembly scheme. MR carried out screening of DNA ligases. PAGT developed the assembly platform and carried out TEM-1 selections. PAGT and VBP conceived the analysis strategy. VBP wrote the analysis algorithm. PAGT carried out the sequence analysis. PAGT and EH carried out the characterization of isolated TEM-1 variants. PAGT and VBP wrote the manuscript.

## Online Methods

### Assembly

All oligos used in InDel assembly were commercially synthesized (Integrated DNA Technologies). Assembly block oligos providing the 5’-end for ligation with the dsDNA template were phosphorylated in 100μl reactions (1 nmol oligo per reaction) containing 1x NEB T4 DNA ligase reaction buffer and 1μl NEB T4 polynucleotide kinase. Reactions were carried out for 3 h at 37°C, followed by inactivation at 80°C for 20 minutes. Oligos were phenol-chloroform extracted, ethanol precipitated, resuspended in 90 μl annealing buffer (10 mM Tris-HCl pH 8.0, 20 mM NaCl, 1 mM MgCl_2_, 0.01% Tween20) and annealed to 1 nmol of the complementary assembly block strand. Building blocks coding for different amino acids were mixed post annealing to create the desired incorporation proportions.

In parallel, 60μl of MyOne C1 streptavidin-coated paramagnetic beads (Thermo Fisher Scientific) were washed twice in BWBS (5 mM Tris-HCl pH7.5, 0.5 mM EDTA, 1 M NaCl, 0.05% Tween20) and incubated at room temperature (in BWBS) for 30 min in a rotating incubator, to reduce background binding. After washing, 10 pmol of biotinylated dsDNA template oligos were added to the beads and incubated overnight at room temperature in a rotating incubator. Beads were washed in BWBS and transferred to a 0.5 ml microcentrifuge tube for assembly.

Bead-bound templates were digested with SapI (NEB) in 100μl reactions (10μl 10x CutSmart buffer, 2μl SapI, 1 μl 1% Tween20) for 2 h at 37°C with vortexing every 15-20 minutes to keep beads in suspension. Beads were isolated and washed once in BWBS. The supernatant containing SapI, was retained and stored at 4°C for subsequent assembly cycles.

The desired mixture of building blocks was added to the washed beads, incubated at 37°C for 30 s, followed by an additional 30 s incubation at 4°C. Supernatant containing the building blocks was removed and beads transferred to a ligation reaction. Ligations were carried out in 100μl reactions (10μl T4 DNA Ligase buffer, 12μl 1,2-propanediol, 10μl 30% PEG-8000, 1μl T4 DNA Ligase, 1μl 1% Tween20, 65μl ddH_2_O) at 25°C for 1 hour, with vortexing every 15-20 minutes.

Beads were isolated, washed in BWBS and could then be taken to start a new assembly cycle. The supernatant containing the ligase reaction mixture was retained and stored at 4°C for subsequent cycles.

The final assembly cycle used a modified dsDNA assembly block (a 3’ cap block) containing a priming site used for post-assembly library amplification. After ligation of the capping oligo, beads were resuspended in 50 μl BWBS for PCR amplification.

### Denaturing polyacrylamide gel electrophoresis

Assembly reactions carried out with FAM-labelled templates could be visualized after separation by denaturing PAGE. Gels were 15% acrylamide (19:1 acrylamide:bis-acrylamide) with 8 M urea in 1x TBE. An equal volume of loading buffer (98% formamide, 10 mM EDTA, 0.02% Orange G) was added to FAM-labelled templates, and sampled were incubated at 95°C for 5 min before being loaded onto the gel. Gels were run at a constant current of 30 mA for 1.5-2 h. FAM-labeled oligos were detected by imaging on a Typhoon FLA 9500 scanner (GE Life Sciences).

### Library amplification and cloning

Assembled libraries were PCR amplified from beads in 50 μl reactions using 10 U MyTaq HS polymerase (Bioline), 0.2 μM each of oligos TEM1-InDel-AmpF(1/2, for corresponding rounds of selection) and TEM1-InDel-AmpR, 1 μl resuspended bead slurry from the assembled library, 1X MyTaq reaction buffer, and 1X CES enhancer solution^31^. Library amplifications were carried out with a 1 min denaturation at 95°C, followed by 20 cycles of 15 s at 95°C, 15s at 55°C, 30 s at 72°C, ending with a 2 min final extension at 72°C. PCR cycles were limited to **20** in library amplifications to minimize amplification biases and reduce likelihood of secondary mutations. Multiple reactions were carried out in parallel to ensure sufficient material for cloning could be generated and the oligos harbored BsaI overhangs for seamless DNA assembly.

Vector backbones were generated by iPCR in 50 μl reactions using 1 U Q5 Hot Start DNA Polymerase (NEB), 0.2 μM each of oligos Vec-TEM1-InDel-F and Vec-TEM1-InDel-R(1/2, for corresponding rounds of selection), 1 ng pTEM1-Cam vector template, 200 μM dNTPs, 1X Q5 reaction buffer, and 1X CES enhancer solution^31^. Vector amplifications were carried out with a 30 s denaturation at 98°C, followed by 30 cycles of 10 s at 98°C, 20 s at **68**°C, and 1.5 mins at 72°C, ending with a 2 min final extension at 72°C. Multiple reactions were carried out in parallel to ensure sufficient material for cloning could be generated and the oligos harbored BsaI overhangs for seamless DNA assembly.

PCR products were purified using NucleoSpin Gel and PCR Cleanup columns (Macherey-Nagel). Purified vector DNA (5 μg) and library (1 μg) DNA were digested with BsaI (NEB) and *Dpn*I (NEB) for 3 h at 37°C in multiple parallel 100 μl reactions and again purified. Vector and library were ligated (1:3 molar ratio, 1 μg total DNA) with NEB T4 DNA ligase for 2 min at 37°C, followed by 6 h at 25°C and overnight at 16°C. DNA was isolated by phenol-chloroform and ethanol-precipitated. Ligated DNA was resuspended in 5μl ddH**2**O and transformed by electroporation into NEB 10-beta cells.

### Selection

Transformed libraries were plated on LB medium supplemented with suitable ceftazidime concentrations for selection, and incubated at 37°C overnight. Colonies were harvested with a cell scraper, transferred to 10 ml LB medium containing ceftazidime, and incubated at 37°C for 2-3 h. The liquid culture was split in three aliquots. One was supplemented with glycerol [to a final 20% (v/v) concentration], and flash frozen for -80°C storage. A second was plated on LB medium containing higher ceftazidime concentrations to isolate the most active TEM-1 variants. The remainder was used for plasmid extraction.

### Antibiotic susceptibility assays

The substrate spectrum of TEM-1 variants was tested by measuring the minimum antibiotic concentration that could inhibit bacterial growth in liquid culture (MIC) and by measuring the growth inhibition of bacteria on solid media. *E. coli* harboring TEM-1 variants were tested for their susceptibility against ampicillin (AMP), carbenicillin (CBN), ceftazidime (CAZ), cefotaxime (CTX) and imipenem (IMP).

For MIC determination, approximately 100 CFU (based on the dilution of a liquid culture in mid-log growth) were added to 200μl LB medium supplemented with different antibiotic concentrations and allowed to grow overnight at 37°C with shaking. MIC assays were carried out in 96-well flat bottom plates (Greiner). Cells were resuspended by mixing with a multichannel pipette and bacterial growth estimated from OD600 measurements. No antibiotic controls were used to estimate the maximum growth of each strain in the experimental conditions and normalize OD600 between independent experiments. Growth inhibition assays in liquid cultures were carried out in triplicate with the lowest concentration of the antibiotic to fully inhibit bacterial growth defining MIC for that strain.

Growth inhibition of the *E. coli* strains in solid medium was carried out using by placing filter paper discs (Oxoid) containing a known amount of each antibiotic onto a lawn of approximately 10^7^ CFU. Antibiotic susceptibility was measured as the radius of growth inhibition around the antibiotic disc. At least three independent experiments were carried out for each strain.

### DNA library preparation for next generation sequencing (NGS)

Libraries for Illumina MiSeq sequencing were prepared by PCR with oligos containing required adaptors and unique indices to allow all pre- and post-selection libraries to be sequenced in a single experiment.

Pre-selection libraries were amplified directly from the streptavidin beads isolated from assembly. Post-selection libraries were amplified from purified plasmid DNA extracted from recovered transformants. Libraries were amplified in 50μl reactions using NEB Q5 Hot-start DNA polymerase to minimize amplification errors and PCR cycles capped at 20 to minimize amplification biases. Reactions contained 1 U polymerase, 0.2μM each of oligos xxx-MiSeqF (separate oligo for each library, with varying index sequences for demultiplexing, names and sequences are in Supplementary Table 2) and TEM1-MiSeq-R, 1 ng plasmid template or 1μl resuspended bead slurry from the assembled library, 200μM dNTPs, 1X Q5 reaction buffer, and 1X CES enhancer solution^31^. Product size and purity were checked on agarose gels and correct amplicons excised and purified using Monarch Gel Extraction (NEB).

Libraries were quantified by fluorimetry using a Qubit 3.0 (Thermo Fisher Scientific) with a dsDNA HS assay kit and pooled proportionally to obtain the desired number of reads for each sample. Sequencing was carried out on an Illumina MiSeq instrument by UCL Genomics using a 150 cycle v3 kit.

### NGS data handling

Sequencing data was treated as described in Supplementary Note 1. Briefly, sequences were filtered for quality, trimmed to keep only the diversified regions, translated into protein sequences, counted, and formatted to serve as input for the *k*-mer analysis.

### NGS analysis

Sequencing was modelled as Poisson distributions, to allow different populations to be compared and enrichment of individual sequences determined. All analyses were carried out in MATLAB (MathWorks). A Z-score, defined in [1] was used as a measure of comparison between pre- and post-selection distributions.

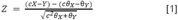

Where c is the ratio in size between post- and pre-selection libraries (to correct for sampling), X is the number of counts for a test sequence in the post-selection library, Y is the number of counts for the same sequence in the pre-selection library. θ_X_ and θ_Y_ are the estimated Poisson parameters (counts as fraction of the total reads) for post- and pre-selection libraries respectively. Z-scores give a measure of enrichment, with extremely positive values identifying the sequences most enriched.

Each sequence was decomposed into all possible masked 3-mers and the library termini encoded as *“Z”* characters (to avoid confusion with natural amino acids and degenerate positions). Masked 3-mers were counted and mapped to a 882-dimension column vector, which each dimension representing one of the possible masked 3-mers. Vectors were normalized and scaled by their Z-score.

Once all sequences identified in selection were assembled in column vectors, primary component analysis (PCA) was carried out to identify dimensions (i.e. masked 3-mers) that contributed the most to selection. Sequence reconstruction was carried out for each of the PCA dimensions using positive components above 0.1 (arbitrarily chosen to minimize noise). Reconstruction was carried out by manual inspection assembling selected sequences from the highest to the lowest PCA coefficient. Reconstruction was successful in most cases generating motifs that encompassed both N- and C-terminal arbitrary “**Ẕ**” characters.

### Loop residue labelling

Numbering of residues within the loop follows the convention for numbering antibody variable regions^23^. Briefly, numbering maintains residues outside the diversified region with their wild-type numbering. Thus, numbering is unchanged for variants of the same length as the wild-type sequence of TEM-1. For variants shorter than wild-type, our scheme introduces a gap between the last diversified codon and downstream invariant position (Table 2).

**Table 1.**
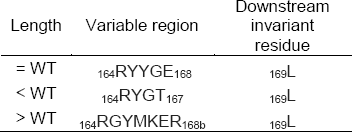
Examples of the proposed numbering scheme for length-variant regions.

Crucially, longer variants are treated as insertions at the end of the diversified region and labelled with an additional lower case letter (e.g. 168a rather than 169) to maintain the residue numbering of the downstream sequence. The proposed numbering scheme unambiguously identifies positions in the sequence without disrupting comparisons between conserved sequence elements outside the library.

